# Interactions of curcumin’s degradation products with the A*β*_42_ dimer: A computational study

**DOI:** 10.1101/2022.08.03.502645

**Authors:** Maryam Haji Dehabadi, Amedeo Caflisch, Ioana M. Ilie, Rohoullah Firouzi

## Abstract

Amyloid-*β* (A*β*) dimers are the smallest toxic species along the amyloid aggregation pathway and among the most-populated oligomeric accumulations present in the brain affected by Alzheimer’s disease (AD). A proposed therapeutic strategy to avoid the aggregation of A*β* into higher order structures is to develop molecules that inhibit the early stages of aggregation, *i*.*e*. dimerization. Under physiological conditions the A*β* dimer is highly dynamic and does not attain a single well defined structure but is rather characterized by an ensemble of conformations. In a recent work, a highly heterogeneous library of conformers of the A*β* dimer was generated by an efficient sampling method with constraints based on ion mobility mass spectrometry data. Here, we make use of the A*β* dimer library to study the interaction with two curcumin degradation products, ferulic aldehyde and vanillin, by molecular dynamics (MD) simulations. Ensemble docking and MD simulations are used to provide atomistic detail of the interactions between the curcumin degradation products and the A*β* dimer. The simulations show that the aromatic residues of A*β*, and in particular ^19^FF^20^ interact with ferulic aldehyde and vanillin through π−π stacking. The binding of these small molecules induces significant changes on the ^16^KLVFF^20^ region.

## 1. INTRODUCTION

The onset of a wide range of neurodegenerative diseases, including Parkinson’s disease (PD), type 2 diabetes, and Alzheimer’s disease (AD) is associated with the aggregation of intrinsically disordered proteins (IDPs) (1,2). AD is the most common neurodegenerative disease currently affecting over 50 million people worldwide and is estimated to reach 131.5 million by 2050 (3-5). The majority of approved drugs against AD treat only the symptoms and fail to prevent or cure the onset of the disease. The pathological hallmark of AD is the presence of extra-cellular senile plaques and intra-cellular neurofibrillary tangles formed from the amyloid-*β* peptide (A*β)* and tau protein, respectively (6). Extra-cellular amyloid fibrils have been proposed to disrupt neuronal communication, while intra-cellular tau-containing tangles can block neuronal transport (2,7-8).

The dominant forms of the A*β* peptide are 40 and 42 residues long, with the latter being the predominant species in amyloid plaques (2). Structurally, the A*β* peptides consist of a disordered N-terminus (first 15 residues), a central hydrophobic cluster (CHC) spanning residues 16-21 and a hydrophobic C-terminus (last 10-12 residues) (9). Amyloid formation follows a nucleation-elongation mechanism (10-11). Briefly, soluble monomers undergo conformational transitions and can aggregate to form dimers, trimers or oligomers, which then evolve to form fibrillar structures (9). Therefore, preventing or reducing the aggregation of the A*β* is an effective therapeutic strategy against AD. In recent years, small molecules have been developed as inhibitors to intervene in the formation of higher order aggregates of A*β* (12-18).

Curcumin is a natural product with anticancer, antioxidant, antiviral, antifungal, anti-inflammatory, and antibacterial properties, which was proposed to have a high therapeutic potential against AD (12,13,17-34). *In vitro* studies showed that curcumin can reduce the *β*- sheet content (24), block A*β* oligomers (24), destabilize and reduce A*β* fibrils (25) and also break the formed tau tangles (26). Furthermore, *in vivo* studies indicated that curcumin can disassemble tau oligomeric species (18,27) and reduce insoluble A*β* oligomers and plaques (28,29), which could limit the progression of AD. Curcumin was shown to have anti-amyloidogenic and fibril-destabilizing effects and to promote the formation of “off-pathway” soluble oligomers and prefibrillar aggregates (25,28,30-31). An *in vivo* study showed that curcumin injected into mice crossed the blood-brain barrier and bound to amyloid plaques (28). Novel curcumin formulations (longvida and theracurmin) administered in low doses (80-180 mg/day) overcome the poor *in vivo* bioavailability (32) of curcumin (33,34). A recent study showed that oral consumption of a bioavailable form of curcumin (theracurmin) led to significant memory improvement, which was attributed to the decrease in amyloid and tau aggregations (34). Moreover, curcumin is unstable in aqueous solution and undergoes rapid hydrolysis (35). Wang et al. found that 90% of curcumin degraded into various products such as ferulic aldehyde, trans-6-(4′-hydroxy-3′-methoxyphenyl)-2,4-dioxo-5-hexenal, feruloyl methane, ferulic acid, and vanillin within 30 min in phosphate buffer at pH 7.4 (35). The degradation products of curcumin have a higher solubility as compared to curcumin and may therefore be appropriate candidate for anti-AD therapeutics (35). As a matter a fact, ferulic acid can protect neurons against A*β*-induced oxidative stress, which plays a significant role in the pathology of AD (36-38). Additionally, ferulic acid can dose-dependently prevent both formation and extension of A*β* oligomers and destabilize A*β* fibrils (39). An *in vitro* study showed that vanillin has acetylcholinesterase inhibitory activities, leading to the restoration of acetylcholine levels in diseased brains leading to an improvement in memory function (40). Vanillin has also been shown to prevent amyloid aggregation *in vivo* (41). Furthermore, vanillin derivatives have exhibited the enhanced antioxidant and anti-acetylcholinesterase properties, which could be as multi-target hybrid compounds for AD treatment (42).

Complementing experiments, computational studies reinforce the fact that curcumin and its products disaggregate and destabilize A*β* protofibrils and fibrils (12,13,43-48), deform the *β*-sheet structure in A*β*_42_ dimers (49) and also form numerous interactions with the A*β* monomer (17,50,51). Salamanova et al. found that curcumin and ferulic acid increase the helical content of the A*β*_42_ peptide (17). A recent computational study consisting of docking followed by multi-ns MD simulations of curcumin and a hexamer peptide model of A*β*_42_ fibril showed that curcumin partly dissociates the tip peptide of the A*β*_42_ fibril by disrupting the *β*-sheet within the ^12^VHHQKLVFF^20^ sequence (43).

Here, we investigate the effects of the curcumin degradation products (ferulic aldehyde and vanillin) on A*β*_42_ dimers by means of computer simulations. In a recent work, we built a diverse conformational library of the A*β*_42_ dimers using the blockwise excursion sampling (BES) and standard CHARMM force fields (52,53). We demonstrated that the conformational library of A*β*_42_ dimers generated by the CHARMM36m force field is in good agreement with experimental data (52). In the present study, we use the generated CHARMM36m library to identify the binding sites of the curcumin degradation products in A*β*_42_ dimers through a new computational pipeline in the framework of the ensemble docking strategy. Next, we perform MD simulations consisting of A*β*_42_ peptide-small molecule complexes to study the effects of the degradation products in a dynamic environment.

## 2. MATERIALS AND METHODS

### 2.1. Preparation of Aβ_42_ dimer library and ligands

The library of A*β*_42_ dimers was generated using the blockwise excursion sampling (BES) protocol, which was recently employed for the construction of a diverse conformational library for A*β*_42_ monomers and dimers (52,53). The BES protocol comprises simulating annealing and many short conventional MD simulations which are called blocks and denoted as **Γ**^***SA***^ and **Γ**^***MD***^, respectively. Two consecutive simulated annealing and MD simulations blocks are referred to as a SA:MD block (**Γ**^***SA*:MD**^), and five **Γ**^***SA*:MD**^ blocks with five different **T**_**max**_ equal to 700 K, 600 K, 500 K, 400 K, and 350 K form an excursion chain (EC). **T**_**max**_ is an important parameter in the simulated annealing block which is the final temperature after the heat-up run. The protocol exploits the ability of simulated annealing to overcome high barriers in the free energy landscape. Each simulated annealing block is followed by a short MD simulation at constant temperature (310 K). The trajectories in MD simulation blocks are used for the sampling of structures. We used 1000 excursion chains (N_EC_ = 1000), and built the initial structures of as extended and parallel A*β*_42_ dimers. For more details on this sampling methodology and its terminology see reference (52,53). The A*β*_42_ dimer structures sampled by the BES protocol were clustered using the Daura algorithm (54) with a Carbon alpha (C*α*) root-mean-square deviation (RMSD) cutoff of 0.3 nm. This analysis produced 41322 clusters. Then collision cross sections (CCS) values were calculated for all structures. A total of 1183 representative structures of all MD snapshots that satisfied the experimental value of CCS (1252 ± 20 Å^2^) were selected (52,55) and used as a library of A*β*_42_ dimers for the docking protocol after energy minimization by 5000 iterations of the steepest descent algorithm. The MD simulations were carried out using the GROMACS 5.1.5 software (56- 57). The CHARMM36m all-atom force field (58) and the Generalized Born (GB) water model (59) were used. The structures of the two degradation products of curcumin (ferulic aldehyde and vanillin, **Fig. 1**) were extracted from the ZINC database (60). The GAMESS package (61) was used to optimize all ligands at the B3LYP exchange and correlation functional (62) and the 6-31+G (d,p) basis set.

**Figure 1.**
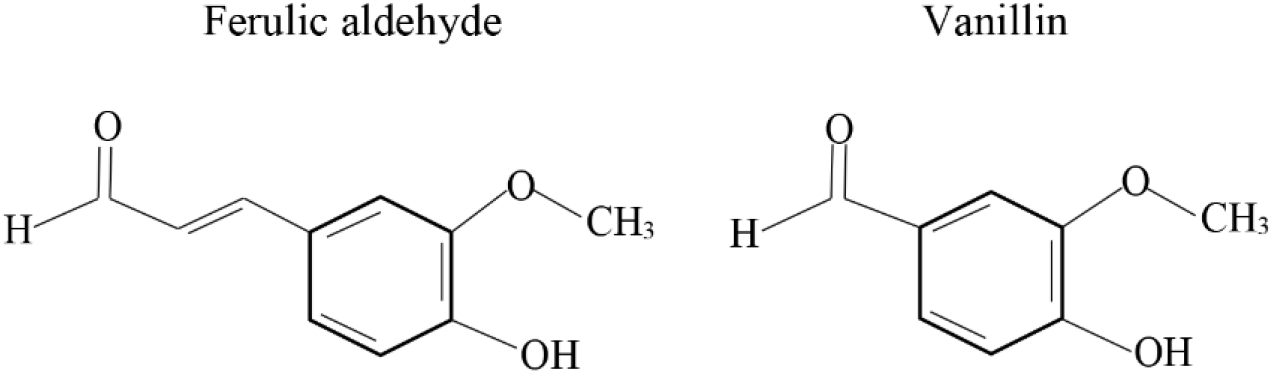
Chemical structure of the degradation products of curcumin used in this study.

### 2.2. Docking setup

The curcumin degradation products were docked against all 1183 A*β*_42_ dimers in the library using AutoDock Vina (version 1.1.2) (63). The docking search space around each representative structure of the A*β*_42_ dimer was defined by a rectangular box centered at the center of mass of the A*β*_42_ dimer. The minimal distance of 1.2 nm from the A*β*_42_ dimer to the edge of box was set. Different optimized docking boxes were defined for each system depending on size and shape of A*β*_42_ dimer conformation. Each docking run produced nine optimal A*β*_42_ dimer–ligand configurations, and totally 10647 (= 1183 × 9) poses were generated. The different poses for each run were ranked by the binding free energy. The top scoring pose for each run is related to the complex with the most favorable binding free energy. The other lower-ranked poses were selected since sometimes the pose with the lowest RMSD relative to the experimental pose is not captured by Vina (14). To select the high-ranked complexes, ***ΔΔG***_***binding***_ was defined as the difference of binding free energy between top-ranked pose and the lower-ranked pose (ΔG_top pose_ ‒ ΔG_lower-ranked pose_). Different cutoff values of ***ΔΔG***_***binding***_ (0.1, 0.2, and 0.3 kcal/mol) were used for selecting the generated docking complexes (**Table 1**). The results were obtained for different cutoff values and for each ligand. The results for ***ΔΔG***_***binding***_ cutoff values of 0.1, 0.2, and 0.3 kcal/mol can be found in the **Supporting Information**.

**Table 1.** The number of selected docking complexes for each ligand with the different ***ΔΔG***_***binding***_ cutoffs.

| $\Delta\Delta G_{binding}$ (kcal/mol) | Ferulic aldehyde | Vanillin |
| --- | --- | --- |
| 0.1 | 1702 | 1679 |
| 0.2 | 2895 | 2746 |
| 0.3 | 4537 | 4240 |

### 2.3 MD simulations setup

#### 2.3.1. System Preparation

Two distinct sets of simulations were set up and simulated following identical protocols. First, ten independent copies consisting of an A*β*_42_ dimer-ligand complex were selected from the highest ranked pose extracted from docking results to investigate the stability of the complex (**Fig. S1** and **S2** in the **Supporting Information**). Second, ten independent copies starting from dimer-ligand complexes, in which the ligand was randomly placed at non-interacting distances from the dimer, were run to identify secondary interaction hotspots on the dimer surface and to investigate the effects of the small molecules on the conformations of the peptides in the presence of each ligand. The initial structure of A*β*_42_ dimer was the most populated structure of A*β*_42_ dimer generated by the BES protocol that has the collision cross sections in agreement with the experimental value. The parameters for all ligands were generated using the CHARMM general force field (CGenFF) (64).

#### 2.3.2 Production Simulations

All simulations were carried out using the GROMACS 2020.4 package (56-57), the CHARMM36m force field (58) and the TIP3P water model (65). Each system was placed in a cubic box of 9 nm per edge and periodic boundary conditions were applied. Explicit Na^+^ and Cl^‒^ ions were added to neutralize the charge of the systems and convey a background concentration of 150 mM. Following the steepest descents minimization, the equilibration was done in the NPT ensemble with position restraints on the backbone of the A*β*_42_ dimer and the heavy atoms of the ligands. The temperature and pressure were maintained constant at 310 K and 1 atm by using the velocity-rescaling thermostat (66) and Berendsen barostat (67), respectively. The first set of simulations, started from the top scoring pose, cumulates 600 ns per ligand (10 independent 60-ns simulations per ligand). The second set of simulations, started from random positions and orientations of the ligands, cumulates 2 µs per ligand (10 independent 200-ns simulations per ligand) in absence of any restraints. All production simulations were started using different initial random velocities and were carried out in the NVT ensemble. The cutoff values for the van der Waals and Coulombic interactions were set to 1.2 nm. The electrostatic interactions were calculated by the particle mesh Ewald (PME) method (68). A time step of 2 fs was employed for all simulations. Bond lengths were constrained using a fourth-order LINCS algorithm (69) with 2 iterations.

## 3. RESULTS AND DISCUSSION

We first investigated the stability of the dimer-ligand complexes starting from the highest ranked docked poses. The analysis revealed that the ligands dissociate from the A*β*_42_ dimer within the first 30 ns. This does not come as a surprise as in the docking step the small molecule is anchored to a rigid interface, while in the MD simulations the complex is highly dynamic (**Fig. S3** and **S4**). Additionally, the conformational changes of the dimer can facilitate formation of binding sites more readily accessible by the ligand. Hence, the reattachment of the small molecule at secondary short-lived interaction sites is observed. Therefore, we performed molecular dynamics simulations starting from non-interacting distances of the small molecules from the A*β*_42_ dimer to eliminate any bias introduced by the initial pose or effects of the rigid dimer conformation. These simulations were used for the analysis. To assess the convergence of docking and MD simulations, the number of contacts between each ligand and individual residues of the two peptides forming the dimer were calculated (**Fig. S5** and **S6** in the **Supporting Information**), and the results indicate that reasonable convergence has been reached.

### 3.1. Identifying the binding site of curcumin compounds

To identify which residues and regions interact most with the ligands, we calculated the average contact number between each A*β*_42_ residue and small-molecule ligand. The results show that the segments ^4^FRHDSGY^10, 19^FF^20, 28^KG^29^, and ^34^LMVGG^38^ interact most with ferulic aldehyde (**Fig. 2a**). Similarly, vanillin interacts predominantly with residue sequences ^9^GY^10, 19^FF^20^, and ^33^GLMVGG^38^ (**Fig. 2b**). Both ligands share the interaction hotspots along the sequence of A*β*_42_ (regions of ^19^FF^20, 34^LMVGG^38^, and ^9^GY^10^), which indicates the homogeneity of the A*β*_42_ dimer molecular recognition sites across the ligands. Importantly, the hotspot regions ^19^FF^20^ and ^34^LMVGG^38^ are part of the central hydrophobic cluster (CHC) and the C-terminus hydrophobic region, respectively, which drive A*β* fibrillization (70-80). In line with the contact number analysis, the analysis obtained from ensemble docking (**Fig. S7-S12** in the **Supporting Information**) indicates that the ligands attach primarily to the CHC region, particularly to its aromatic residues (F19 and F20). These observations are consistent with previous molecular dynamics simulations, which showed that curcumin forms longer lived contacts with residues I32, L34, and M35, while the aromatic residues F19 and F20 showed a high propensity to interact with the rings of curcumin (49). Based on the fragment mapping calculations on the monomeric form of the A*β*_42_ peptide, Zhu et al. reported F4 and Y10 together with L17, F19, I31, and M35 as binding “hotspots” of curcumin in the A*β*_42_ peptide (51). Moreover, solid-state NMR experiments suggested that the CHC segment is a possible binding site for curcumin (43,48,81). Taken together, these results show that the curcumin compounds have a high propensity to interact with the CHC and, in particular, with its aromatic residues. Importantly, the ^17^LVFF^20^ segment has been shown to play a central role in A*β* misfolding and aggregation (70-76,80).

**Figure 2.**
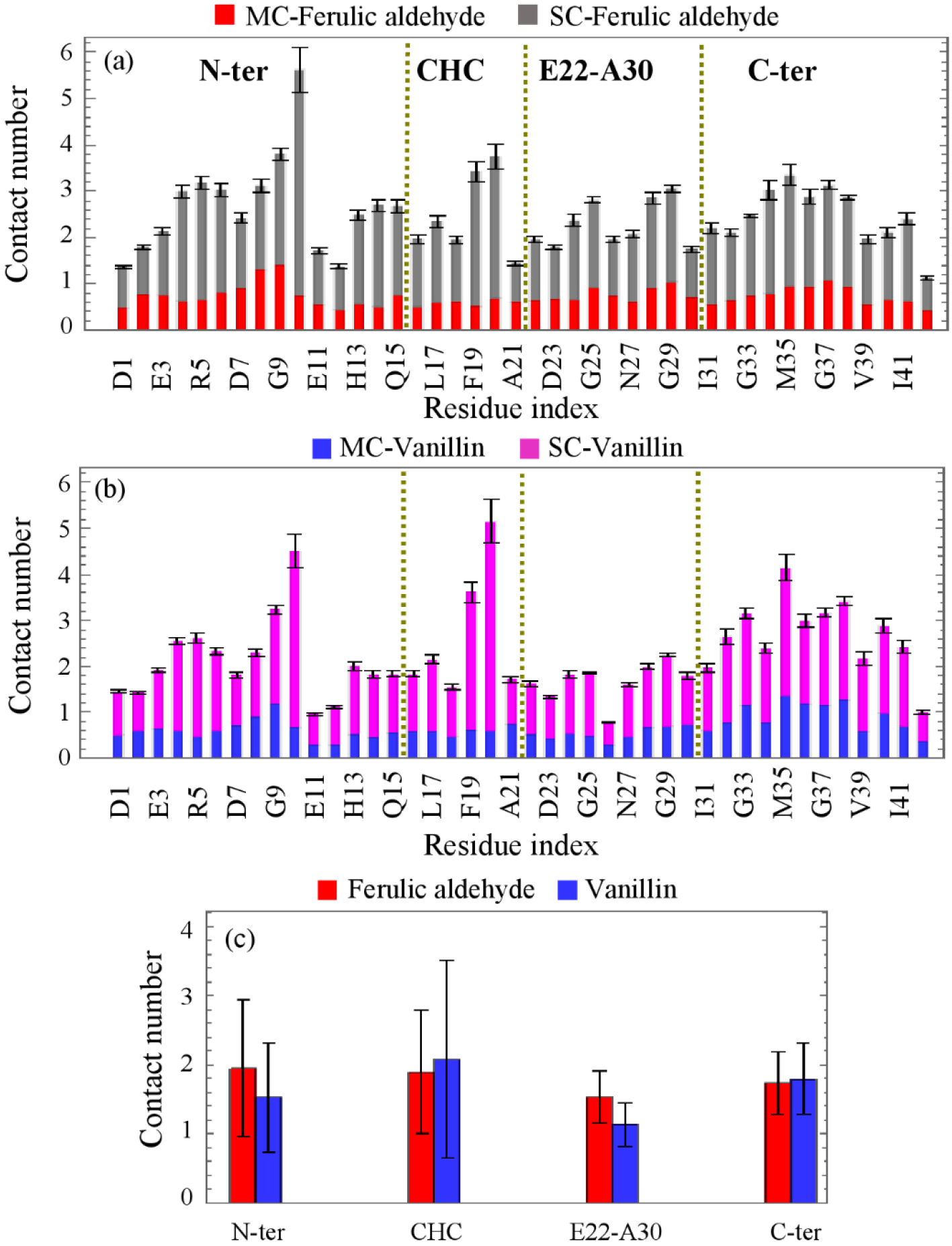
The average number of contacts between main-chain (MC)/ side-chain (SC) of each residue in A*β*_42_ dimer and (a) ferulic aldehyde, (b) vanillin. (c) The averaged contact number for four different regions of A*β* peptide; N-terminus (N-ter), CHC, ^22^EDVGSNKGA^30^ (E22-A30), and C-terminus (C-ter). The error bars represent the standard deviation of the mean. For figure 2a-b, the error bars correspond to the standard deviation of the mean of the cumulated MC and SC contacts. The dashed lines separate the different regions of A*β* (N-ter, CHC, E22-A30, and C-ter). The number of contacts were calculated for the cutoff distance of 0.5 nm.

The contact map analysis reveals that the phenyl ring and the methoxy group (OCH3) in ferulic aldehyde interact more frequently with the residues in CHC region than vanillin (**Fig. 3a-b**). Additionally, all groups of the ferulic aldehyde show a high contact probability with the N-terminus segment compared to these groups in vanillin, while in presence of vanillin, all groups form more interactions with residues in the C-terminus region than ferulic aldehyde, hence could be a main factor in increase of the averaged contact numbers with this region (see **Fig. 2c and Fig. 3b**). The hydroxyl group (OH), methoxy group (OCH3), and aldehyde group (CHO) in ferulic aldehyde and vanillin can interact with the A*β* dimer through the formation of hydrogen bonds and π−π stacking with the A*β*_42_ dimer.

**Figure 3.**
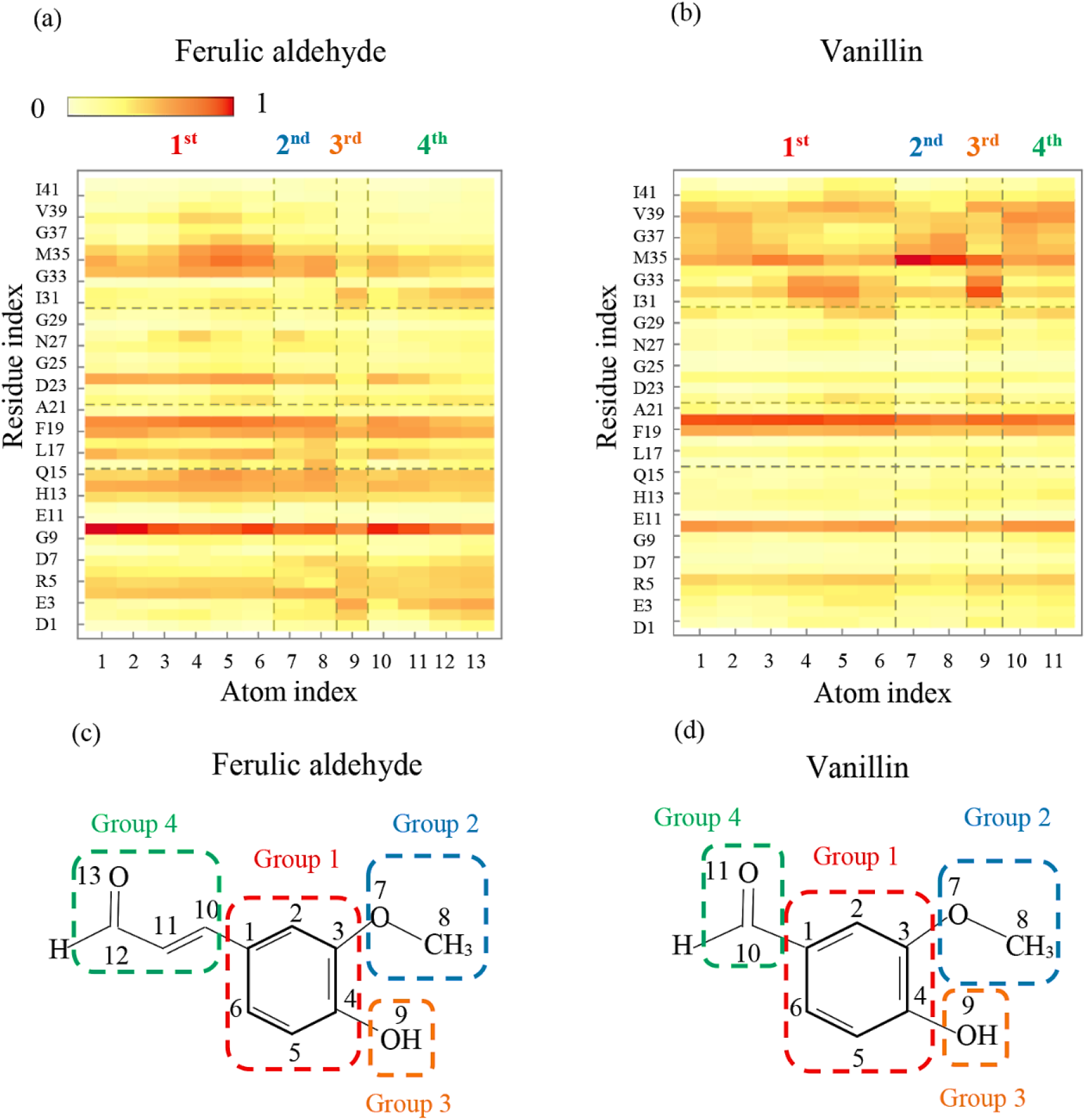
Contact map between the heavy atoms of A*β*_42_ residues and (a) ferulic aldehyde, (b) vanillin. (c-d) Structure of each ligand decomposed into four functional groups. The horizontal and vertical dashes correspond to the different regions of A*β* (N-terminus, CHC, ^22^EDVGSNKGA^30^, and C-terminus) and functional groups of each ligand, respectively. A contact is formed when the distance between the heavy atoms is less than 0.5 nm.

### 3.2. Effective interactions in Aβ_42_ dimer-ligand complex

The ligand-peptide hydrogen bond (H-bond) analysis shows that ferulic aldehyde forms the more frequent of H-bonds with residues Q15 and L34 than vanillin (**Fig. 4a-b**). A similar observation was drawn by Zhao et al. in studying the effect of curcumin on the stability of A*β* dimers in which it was shown that curcumin predominantly interacts with residues Q15 and L34 (49). Apart from curcumin, epigallocatechin (EGC) exhibited high hydrogen bonding preference with residue Q15 (15). Compared to ferulic aldehyde, vanillin forms more H-bonds with the ^31^IIG^33^ and ^40^VI^41^ sequences in the C-terminus region (**Fig. 4c**). In fact, the strong interaction of vanillin with the C-terminal residues of the A*β*_42_ peptides results from the increased hydrogen bond contacts formed by its hydroxyl group (OH), methoxy group (OCH3), and aldehyde group (CHO) with residues G33, V40, and I41.

**Figure 4.**
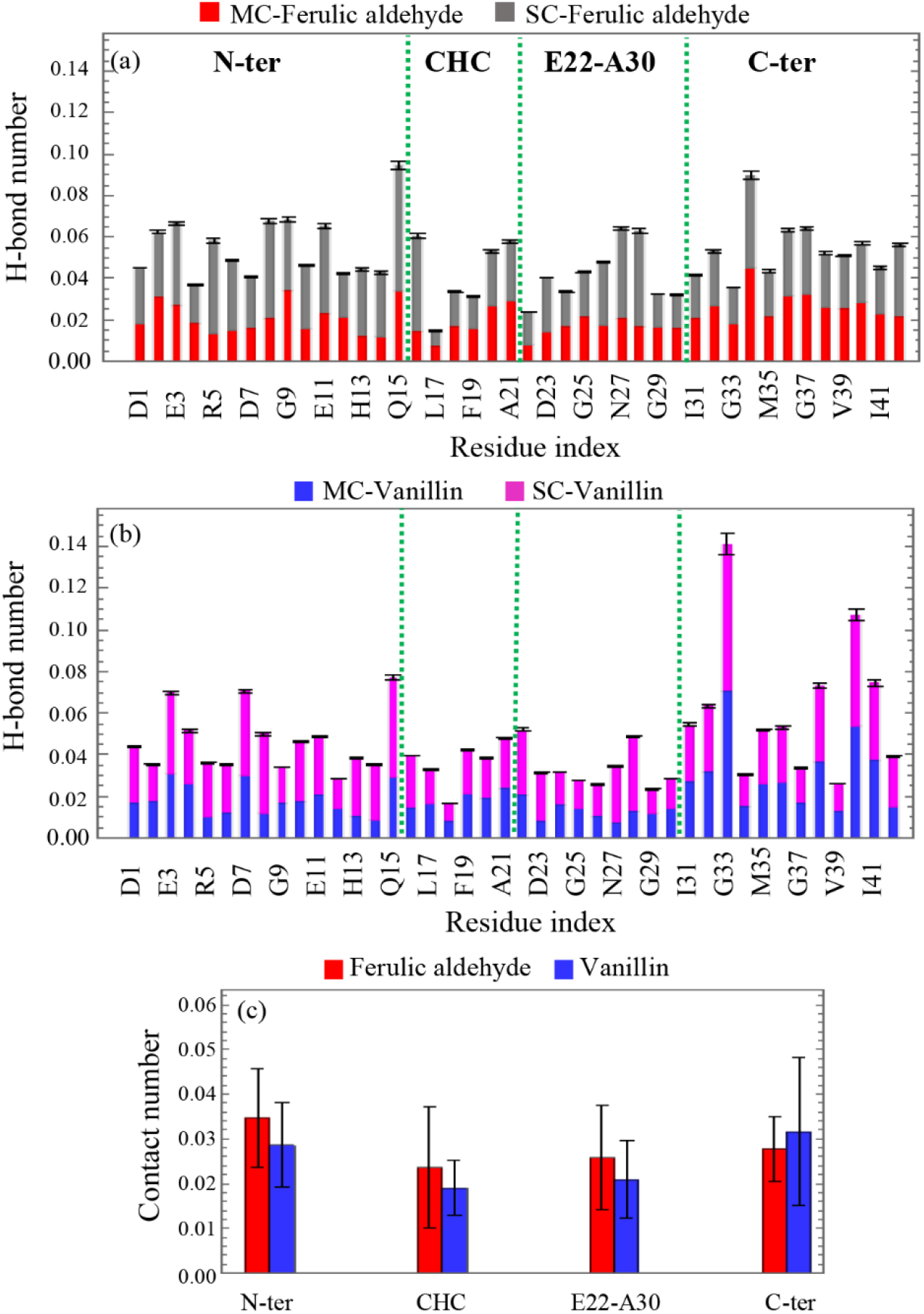
The average H-bond number between main-chain (MC)/ side-chain (SC) of each residue in A*β*_42_ dimer and (a) ferulic aldehyde, (b) vanillin. (c) The averaged H-bond number for four different regions of A*β* peptide; N-terminus (N-ter), CHC, ^22^EDVGSNKGA^30^ (E22-A30), and C-terminus (C-ter). A H-bond is formed if the acceptor-donor distance and acceptor-donor-hydrogen angle are less than 0.35 nm and 30 respectively (15). The error bars represent the standard deviation of the mean. The green dashed lines indicate the different regions of A*β* peptide; N-terminus (N-ter), CHC, ^22^EDVGSNKGA^30^ (E22-A30), and C-terminus (C-ter).

Both ferulic aldehyde and vanillin interact with high propensity with residues Y10, F19, and F20 via π−π stacking (**Fig. 5a**). These observations are in line with the previous computational studies reporting that curcumin interacts with A*β*_42_ via π−π stacking with the side chains of F4, Y10, F19, and F20 in monomers and also H14 for A*β* in dimers and fibrils (13,43,44,46,49,51). Furthermore, we examined the stacking pattern between each ligand ring and each aromatic residue by calculating the centroid distance and angle between the two rings (**Fig. 5b-d**). Ferulic aldehyde prefers to form parallel π−π stacking with Y10 and T-shaped π−π stacking with F20 (**Fig. 5e-f**). Vanillin forms T-shaped π−π stacking with Y10 and parallel π−π stacking with F20 (**Fig. 5g**). **Fig. 5b-d** show that the parallel and T-shaped stacking patterns are the dominant packing modes for the aromatic residues with a high number of π−π stacking. From the H-bond and π−π stacking analysis (**Fig. 4 and Fig. 5**), we identify that ferulic aldehyde more effectively interacts with the dimer by the increasing the number of H-bonds formed by its hydroxyl group (OH), methoxy group (OCH3), and aldehyde group (CHO) with the CHC and N-terminus segments and more π−π stacking interactions with F4, H6, Y10, and H14 as compared to vanillin.

**Figure 5.**
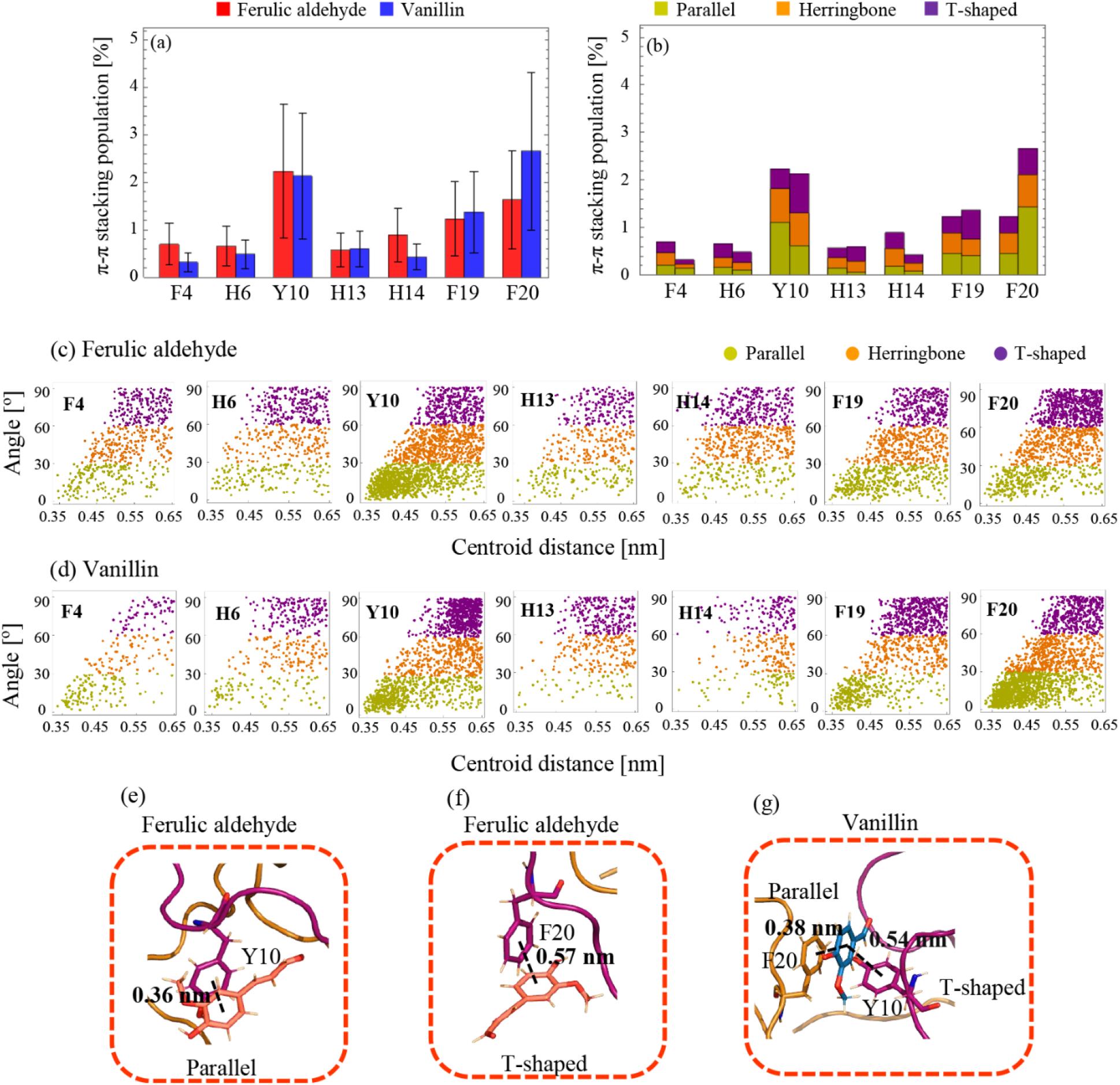
(a) The population (in %) of π−π stacking between the phenyl ring of each ligand and aromatic residue within A*β*_42_ dimer. (b) The population (in %) of parallel, herringbone, and T-shaped stacking between the phenyl ring of each ligand and aromatic residue. Distributions of the centroid distance and angle between two phenyl rings of each aromatic residue and ligand; (c) Ferulic aldehyde, (d) Vanillin. The π−π interaction is formed if the centroid distance between two phenyl rings is less than 0.65 nm. The π−π stacking patterns are classified into three categories: parallel (0-30֯), herringbone (30-60֯), and perpendicular or T-shaped (60-90֯) (15). The π−π stacking interactions were calculated by codes written in Mathematica software. Representative stacking patterns showing the parallel π−π stacking between Y10 and ferulic aldehyde and T-shaped stacking pattern between F20 and ferulic aldehyde (e-f), T-shaped stacking between Y10 and vanillin and parallel stacking pattern between F20 and vanillin (g). The interacting aromatic residues in chainA and chainB are colored in purple and brown, respectively. π−π interactions are given by black dashes. Ferulic aldehyde and vanillin are represented in stick with two colors; pink and blue, respectively.

### 3.3. The disruptive and inhibitor effects of curcumin products on the Aβ_42_ dimer

The average C_*α*_-RMSD values for the full length A*β*_42_ dimer and of four individual segments (N-terminus, CHC, ^22^EDVGSNKGA^30^ and C-terminus) show no significant differences in presence of the small molecules (**Fig. 6a)**. To further characterize the effects of the degradation products of curcumin, we calculated the intra-molecular salt bridge populations (**Fig. 6b**). First, we analyzed the salt-bridges identified from the known fibrillar structures of the peptide. The most recent cryo-electron microscopy structure of the full-length A*β*_42_ fibril (LS-shape fibril structure; PDB id: 5OQV (82)) identified the intra-molecular salt bridges D7-R5, E11-H6 and E11-H13, and K28-A42 (82). Furthermore, the K28-A42 salt bridge is present in the S-shaped A*β*_11-42_ (PDB id: 2MXU) (83) and A*β*_15-42_ (PDB id: 5KK3) (84) and also LS-shaped A*β*_42_ fibrils. Our simulations reveal that the D7-R5, E11-H6, E11-H13 and K28-A42 salt bridges are only sporadically populated in presence of the small molecules. On the other hand, we find that the small molecules have a more pronounced effect on other salt-bridges. For instance, ferulic aldehyde stimulates the formation of the D7-K28, E11-K28 and E22-K28 salt-bridges, while vanillin impacts predominantly the D7-K16. D7-K28, E11-H13, E11-K28 and D23-K28 contacts. Taken together, it is evident that the ligands have a different impact on the formation of the E22-K28 and D23-K28 salt-bridges.

**Figure 6.**
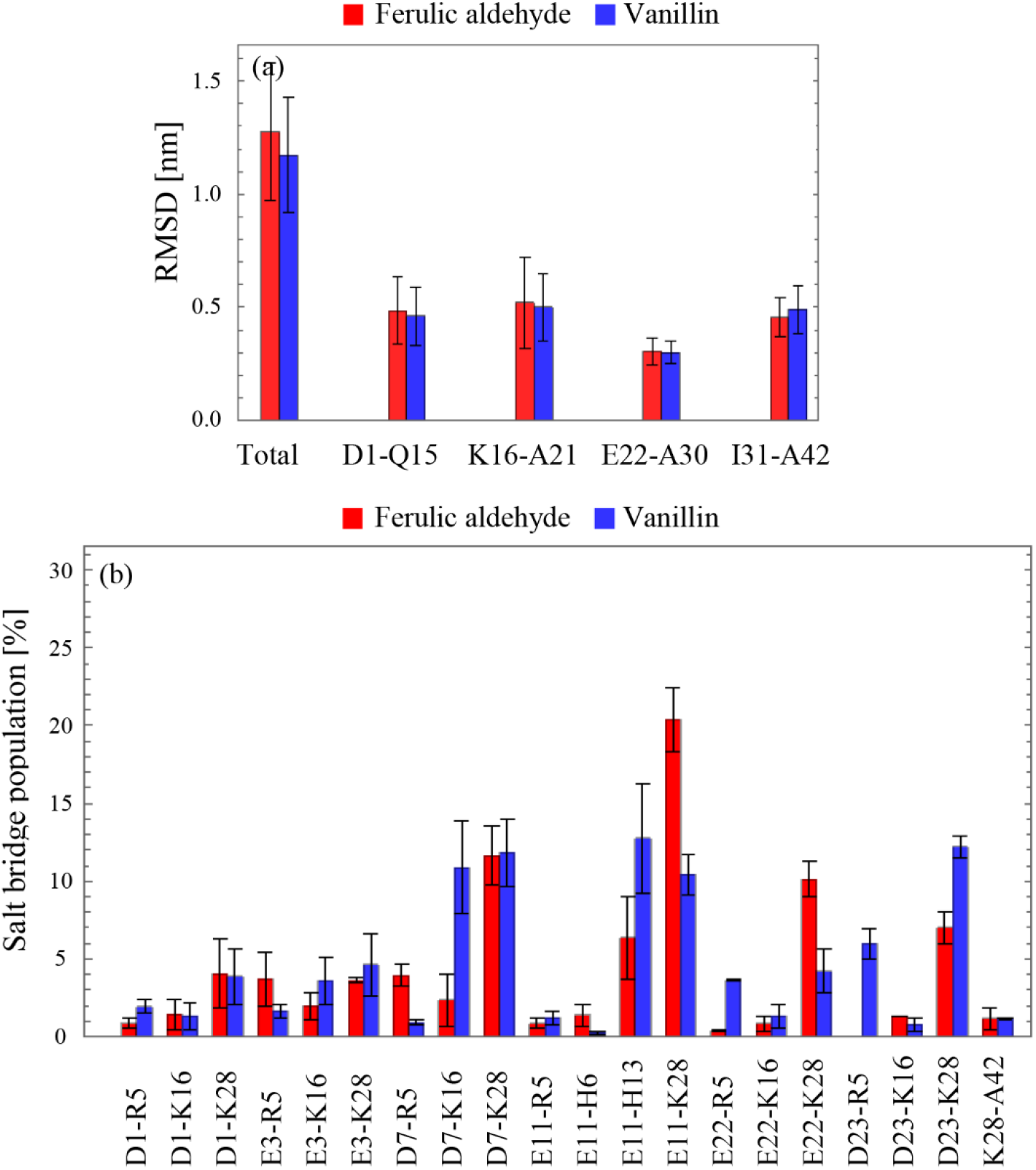
(a) The averaged and standard deviation of C*α*-RMSD for the total and four regions (N-ter, CHC, E22-A30, and C-ter) in A*β*_42_ peptide in the presence of ferulic aldehyde and vanillin. (b) The population (in %) of intra-molecular salt bridges formed in A*β*_42_ dimer in the presence of ferulic aldehyde and vanillin as a ligand. The Cα-RMSD of sampled A*β*_42_ dimers were analyzed by GROMACS tools. The RMSD was calculated for the Carbon alpha (C*α*) as 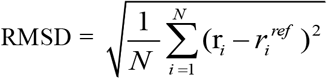 where 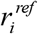 represents the reference position of atom i, and N shows the number of carbon atoms belonging to total and each segment in dimer. The error bars represent the standard deviation of the mean. A salt bridge is formed when the minimum distance between the charged atoms is equal or less than 0.35 nm (85).

## 4. CONCLUSIONS

Inhibition of the A*β* dimerization is a challenging task due to the intrinsic plasticity of the A*β*_42_ dimers. The structural heterogeneity and transient kinetics are a major obstacle for theoretical and experimental methods. To tackle this problem, several sampling methods have been proposed for constructing a representative conformational ensemble for A*β*. To this end, we have very recently employed the BES method for sampling of the A*β*_42_ dimers and shown that BES generated a heterogeneous conformational library of the A*β*_42_ dimers in good agreement with experimental data (52). Here, we investigated the interaction between the A*β*_42_ dimers sampled by the BES protocol and two curcumin degradation products, ferulic aldehyde and vanillin, by ensemble docking and MD simulations (cumulative sampling of 2 µs).

Overall, the simulation results indicate that the curcumin products share the same binding “hot spot” (^16^KLVFFA^21^) on the A*β*_42_ dimer. Importantly, the interaction between the ^16^KLVFFA^21^ regions in A*β*_42_ monomers is key for fibril formation and stabilization (70-76). The contact number analysis provides evidence that ferulic aldehyde interacts with the A*β*_42_ dimer more effectively than vanillin. Ferulic aldehyde preferentially interacts with the N-terminus residues F4, Y10, H14, and also residue F19 and F20 in the CHC segment, whereas vanillin interacts with the A*β*_42_ dimer via π−π stacking interactions with residue Y10, F19, and F20. Moreover, vanillin forms more hydrogen bond interactions with the C-terminus residues in the A*β*_42_ dimer than ferulic aldehyde.

The RMSD values calculated for each region of the A*β*_42_ dimer show that both ligands mainly affect the CHC region without dissociating the dimer. In presence of ferulic aldehyde, one observes the small population of the intra-molecular salt bridges E11-H6 and K28-A42 that are important to stabilize the LS shape of fibril structure. Furthermore, the E11-H13 salt bridge is more stable in the presence of vanillin. On the basis of these results, ferulic aldehyde is predicted to be a more effective inhibitor of A*β*_42_ dimerization than vanillin. *in vivo* data shows that ferulic aldehyde effectively reduces the A*β* deposits and the toxic soluble A*β* oligomers, thus eliminating AD-like pathological changes in the hippocampus and cerebral cortex and improving learning and memory capacity of the model mice (86). Furthermore, both *in vitro* and *in vivo* studies show that vanillin and its derivatives have inhibitory potential against acetylcholinesterase, anti-oxidant properties, and amyloid aggregation inhibitory effects (40-42,87,88).

In conclusion, the degradation compounds of curcumin can inhibit the A*β* association by effective interactions with the aromatic residues ^19^FF^20^ in the CHC region. The simulation results also reveal the presence of hydrogen bonds between the OH, OCH3, and CHO groups on the phenyl rings of the degradation products and residues in the N-terminus and C-terminus of A*β*. This work sheds light on the influence of curcumin degradation products on the A*β* dimer, which may be beneficial for the design of drug candidates.

## AUTHOR CONTRIBUTIONS

A.C., I.M.I. and R.F. designed and directed the project. R.F. performed and analyzed the docking simulations. M.H.D performed and analyzed the MD simulations. M.H.D prepared the manuscript. All authors discussed the results and reviewed the manuscript.

## ACKNOWLEDGMENTS

The computational resources were provided by the Swiss National Supercomputing Centre (CSCS) in Lugano.

## Table of Contents

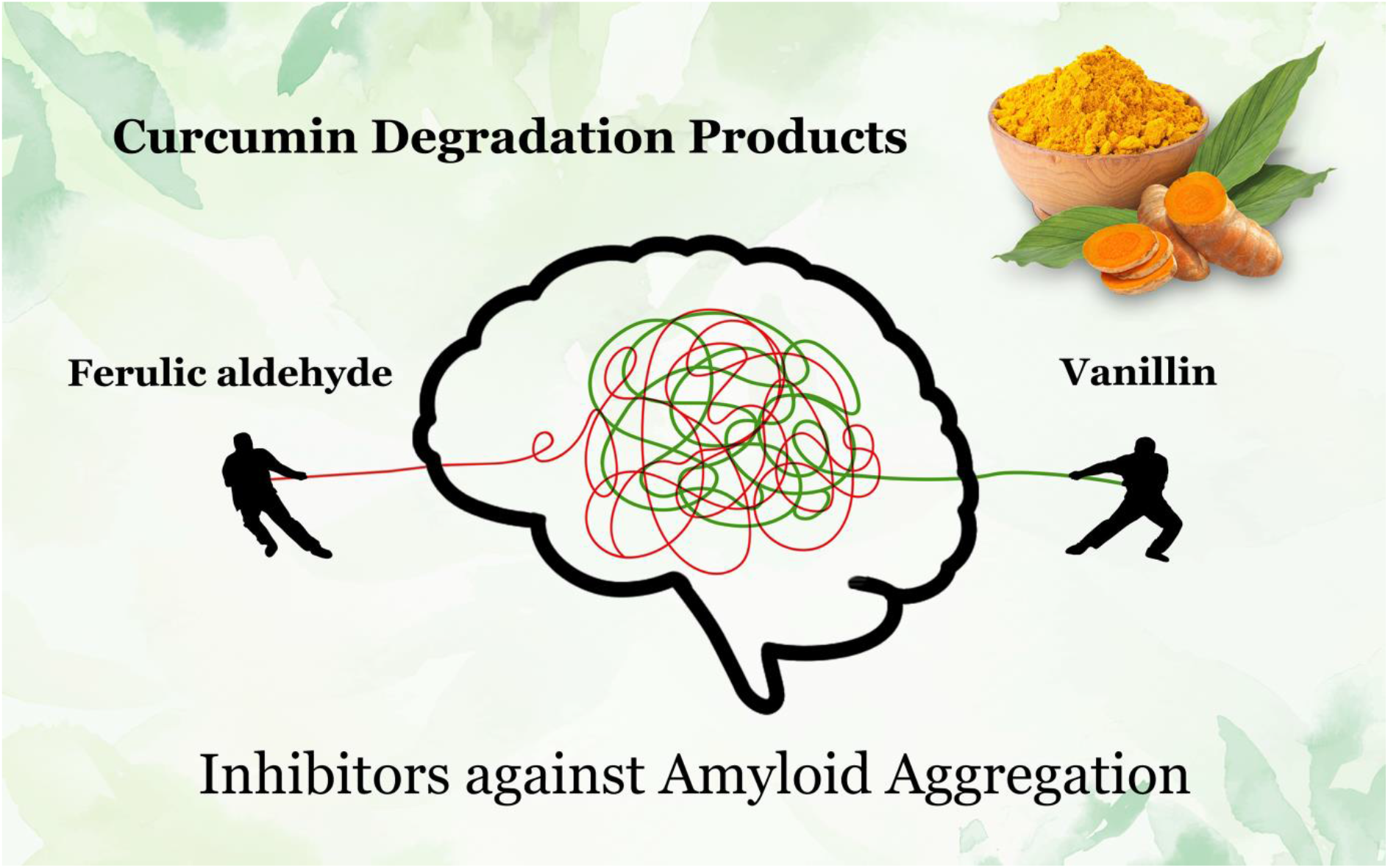

