## Supporting Information for "Interactions of curcumin’s degradation products with the A*β*_42_ dimer: A computational study"

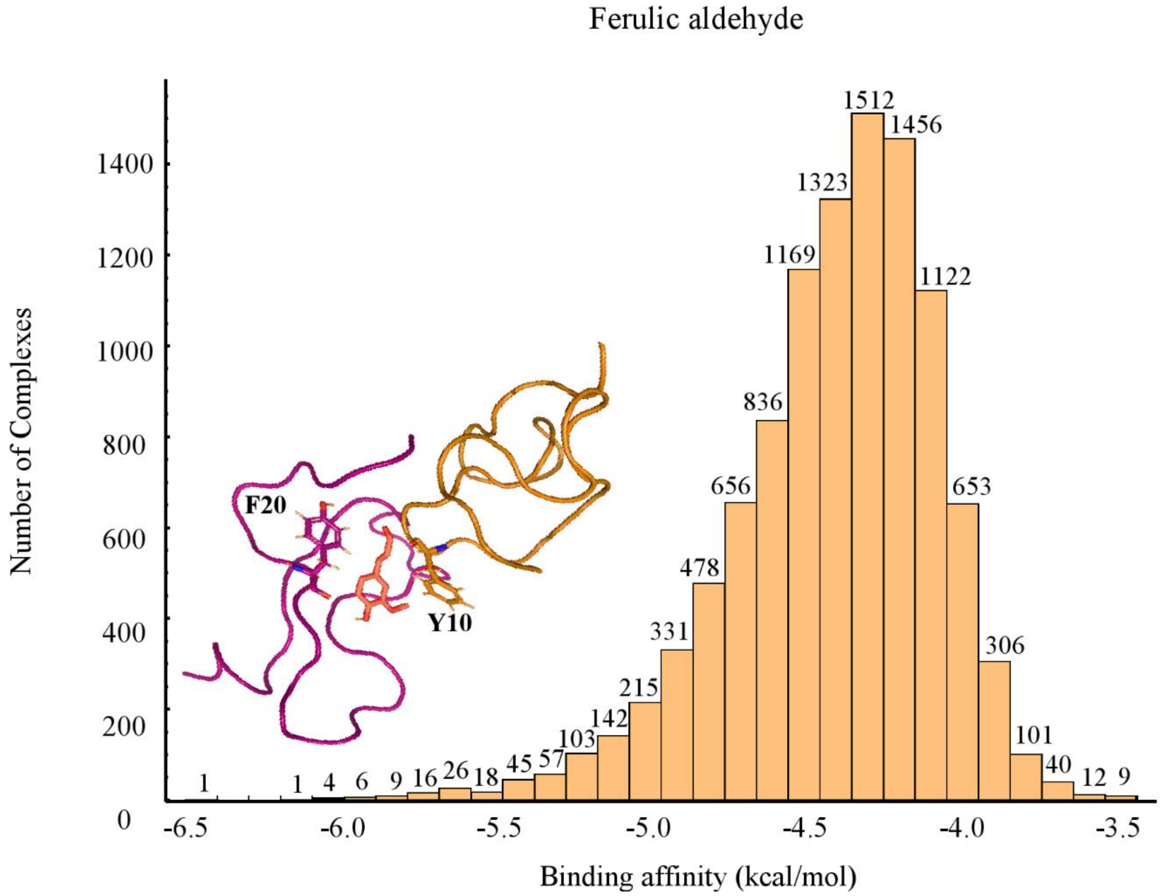


**Figure S1.** The histogram of binding affinity (kcal/mol) of ferulic aldehyde over the $\Delta\Delta G_{binding}$. The binding affinities were divided to bins of 0.1 kcal/mol, and the numbers on top of the bins show the number of complexes for each bin. The interacting aromatic residues with the ligand for some complexes with the highest binding affinities are shown in licorice representation. The chainA and chainB are colored in purple and brown, respectively. Ferulic aldehyde is represented in stick with pink color.

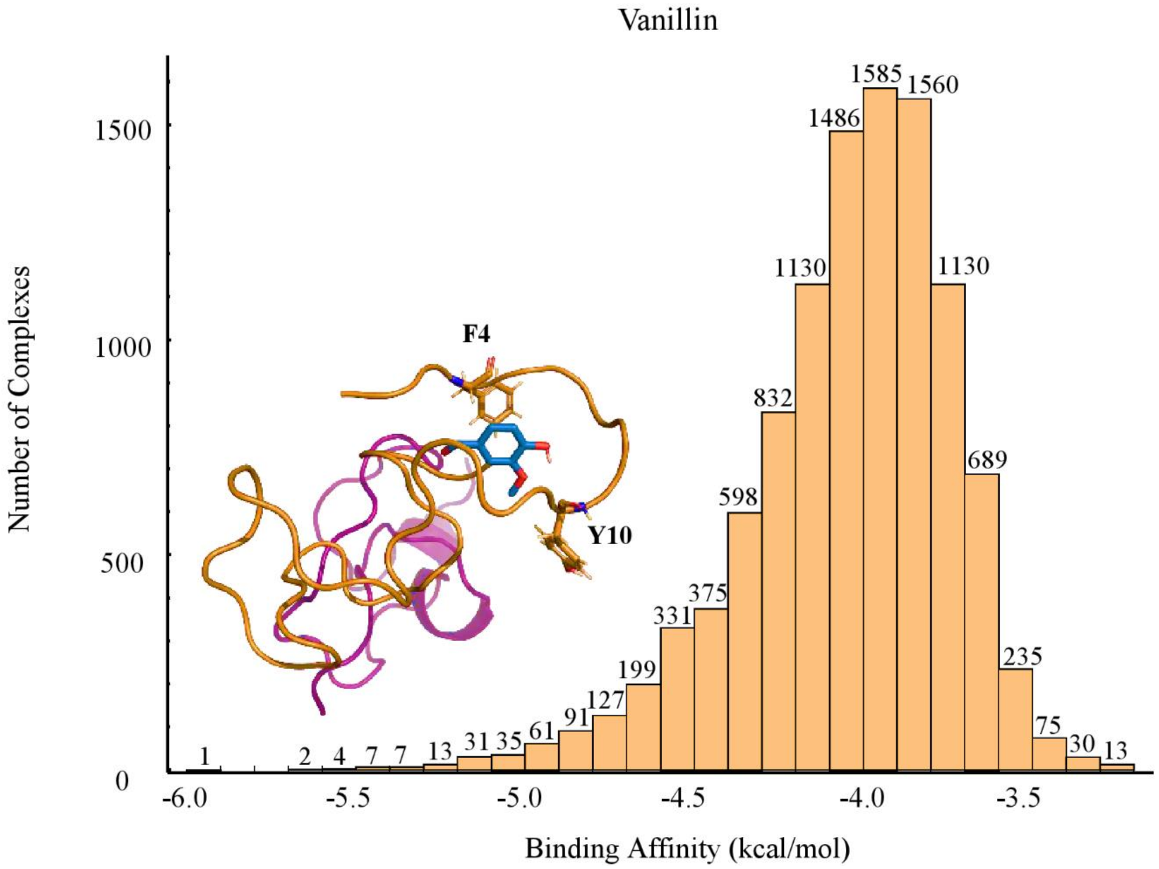


**Figure S2.** The histogram of binding affinity (kcal/mol) of vanillin over the $\Delta\Delta G_{binding}$. The binding affinities were divided to bins of 0.1 kcal/mol, and the numbers on top of the bins show the number of complexes for each bin. The interacting aromatic residues with the ligand for some complexes with the highest binding affinities are shown in licorice representation. The chainA and chainB are colored in purple and brown, respectively. Vanillin is represented in stick with blue color.

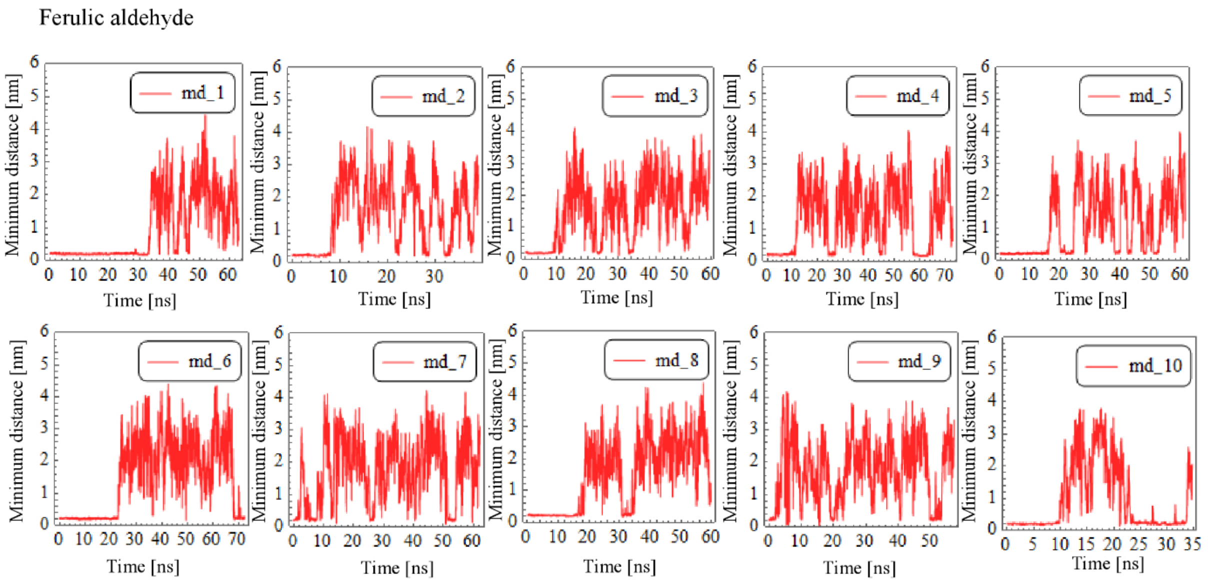


**Figure S3.** The minimum distance between ferulic aldehyde and A$\beta_{42}$ dimer structure with the highest binding affinity obtained by docking simulations as the initial structure.

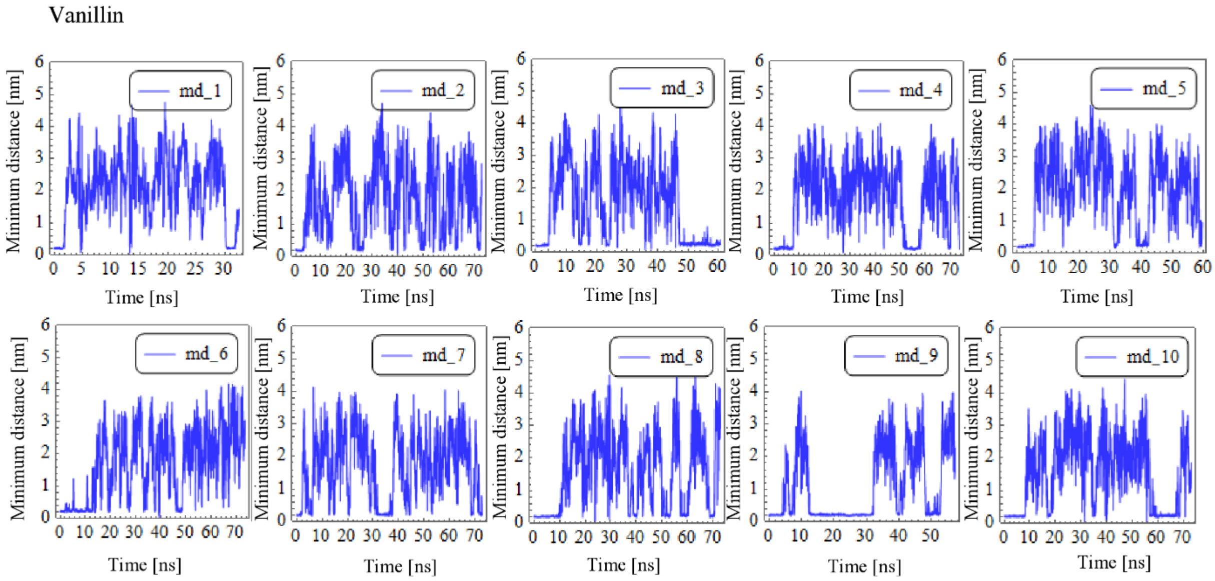


**Figure S4.** The minimum distance between vanillin and A$\beta_{42}$ dimer structure with the highest binding affinity obtained by docking simulations as the initial structure.

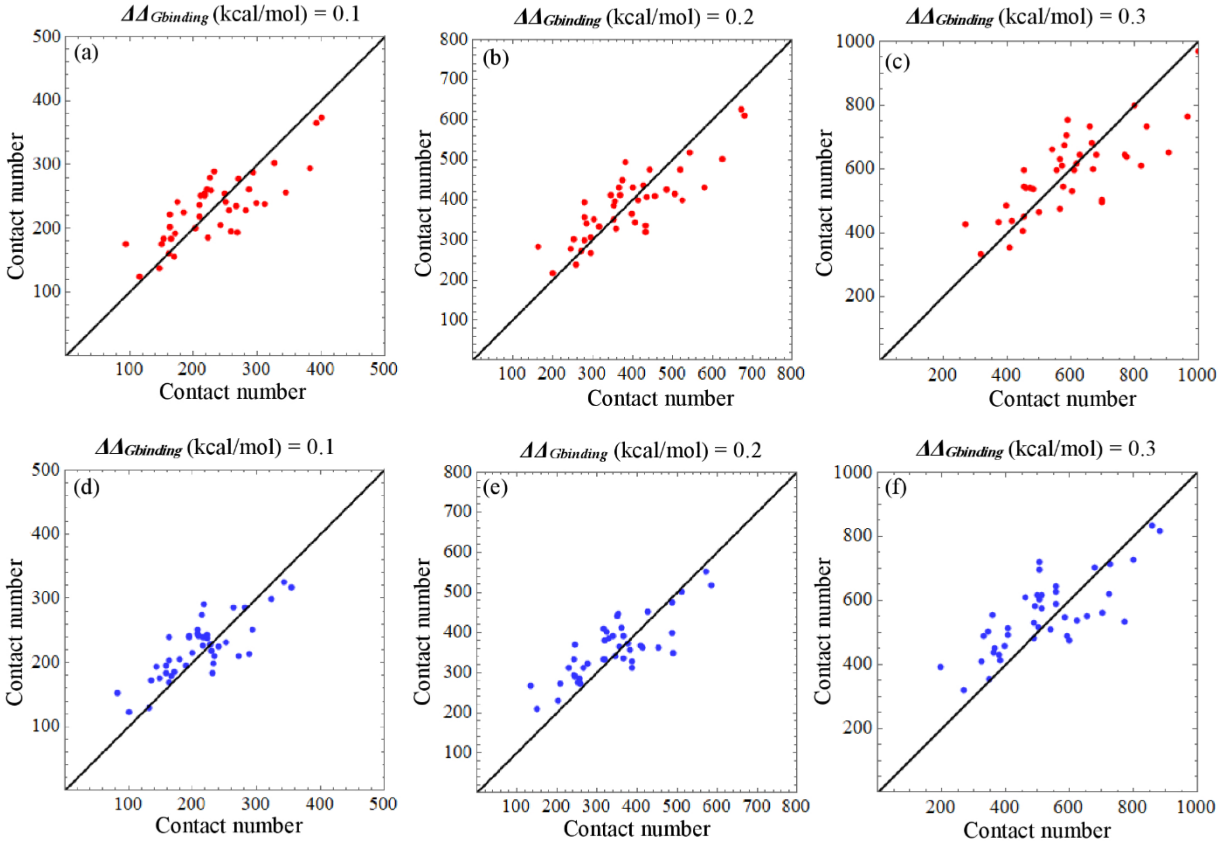


**Figure S5**. The number of contacts between each ligand and individual residues of A$\beta_{42}$ dimer sampled by docking simulations. (a-c) Convergence between the number of contacts of each residue in the first and second half of the sequence and ferulic aldehyde for different $\Delta\Delta G_{\text{binding}}$. (d-f) Convergence between the number of contacts of each residue in the first and second half of the sequence and vanillin for different $\Delta\Delta G_{\text{binding}}$. The horizontal and vertical axes show the contact number for chainA and chainB, respectively. The number of contacts were calculated for the cutoff distance of 0.5 nm in all plots.

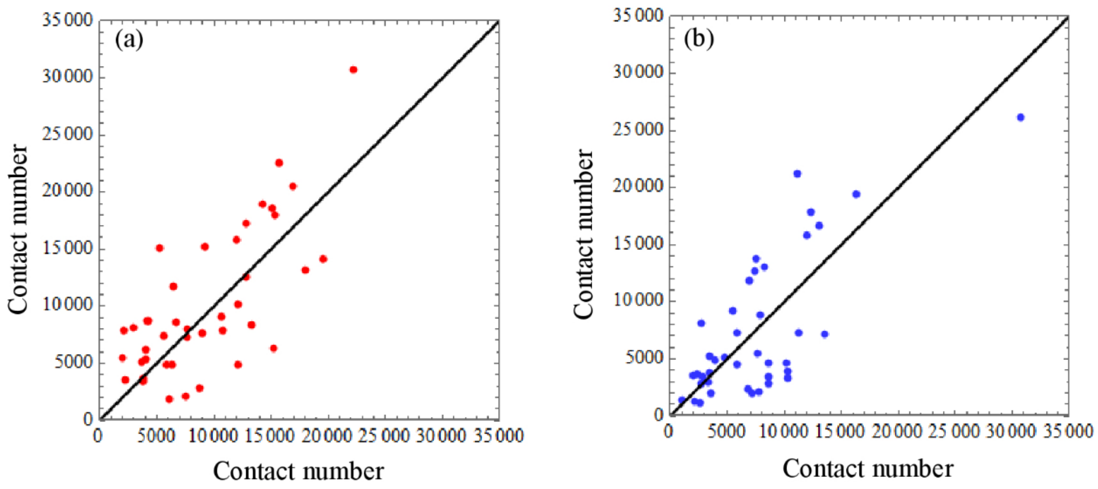


**Figure S6**. The number of contacts between each ligand and individual residues of A$\beta_{42}$ dimer sampled by MD simulations. (a) Convergence between the number of contacts of each residue in the first and second half of the sequence and ferulic aldehyde. (b) Convergence between the number of contacts of each residue in the first and second half of the sequence and vanillin. The horizontal and vertical axes show the contact number for chainA and chainB, respectively. The number of contacts were calculated for the cutoff distance of 0.5 nm in all plots.

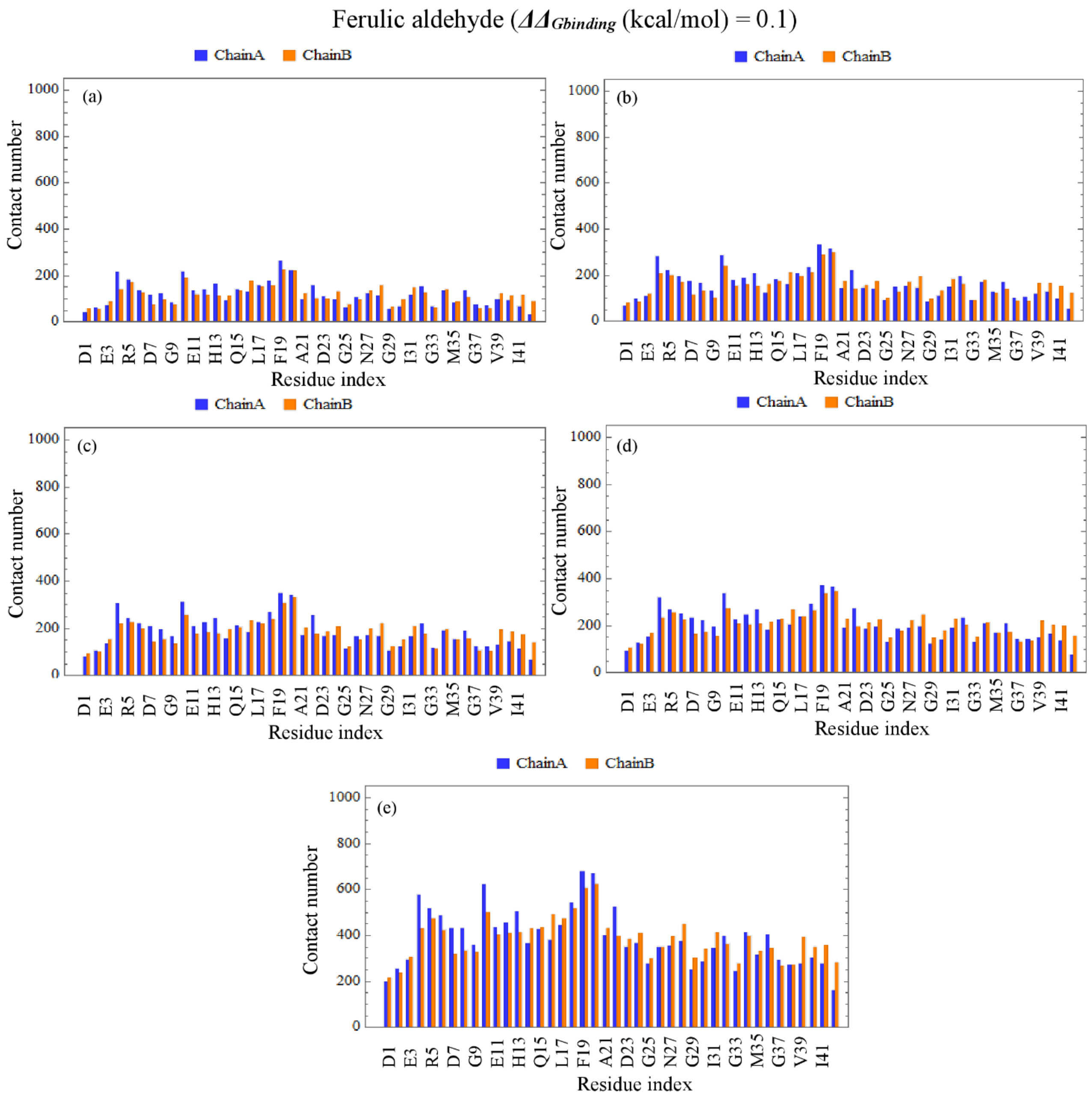


**Figure S7.** The number of contacts between individual residues and ferulic aldehyde for $\Delta\Delta G_{binding}$ (kcal/mol) = 0.1 with five different distance cutoffs; (a) 0.3 nm, (b) 0.35 nm, (c) 0.4 nm, (d) 0.45 nm, and (e) 0.5 nm.

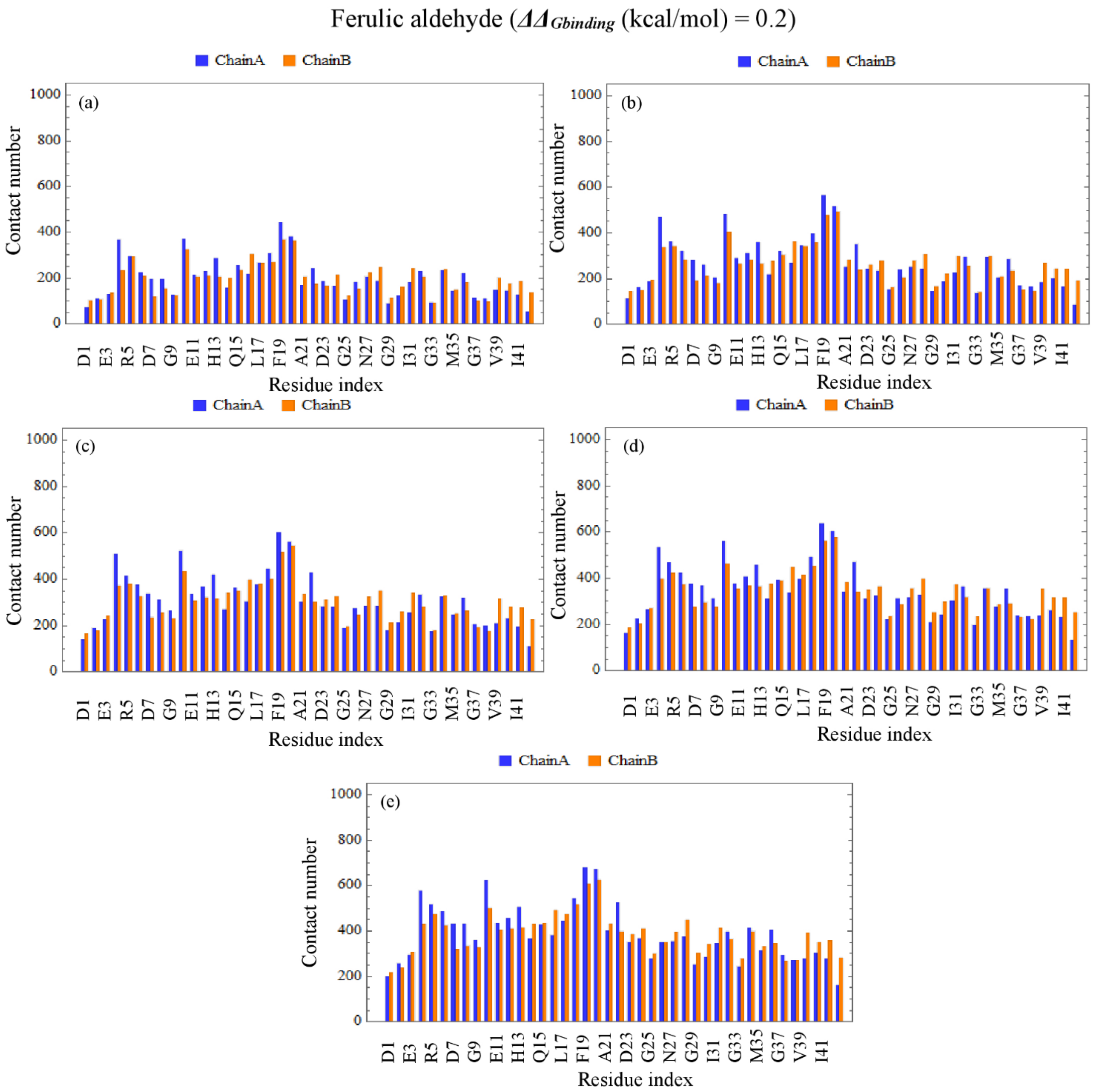


**Figure S8.** The number of contacts between individual residues and ferulic aldehyde for $\Delta\Delta G_{binding}$ (kcal/mol) = 0.2 with five different distance cutoffs; (a) 0.3 nm, (b) 0.35 nm, (c) 0.4 nm, (d) 0.45 nm, and (e) 0.5 nm.

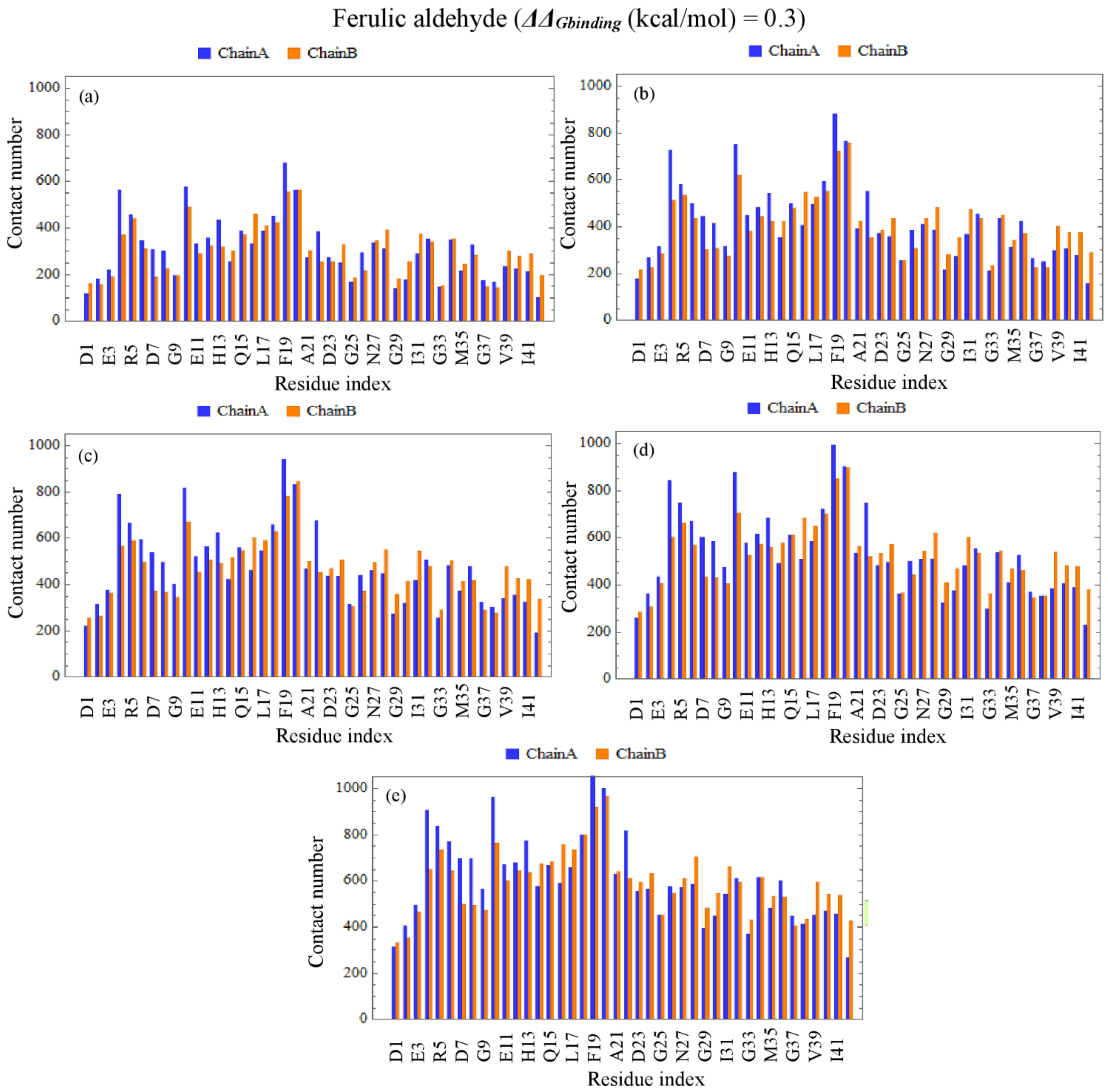


**Figure S9.** The number of contacts between individual residues and ferulic aldehyde for $\Delta\Delta G_{binding}$ (kcal/mol) = 0.3 with five different distance cutoffs; (a) 0.3 nm, (b) 0.35 nm, (c) 0.4 nm, (d) 0.45 nm, and (e) 0.5 nm.

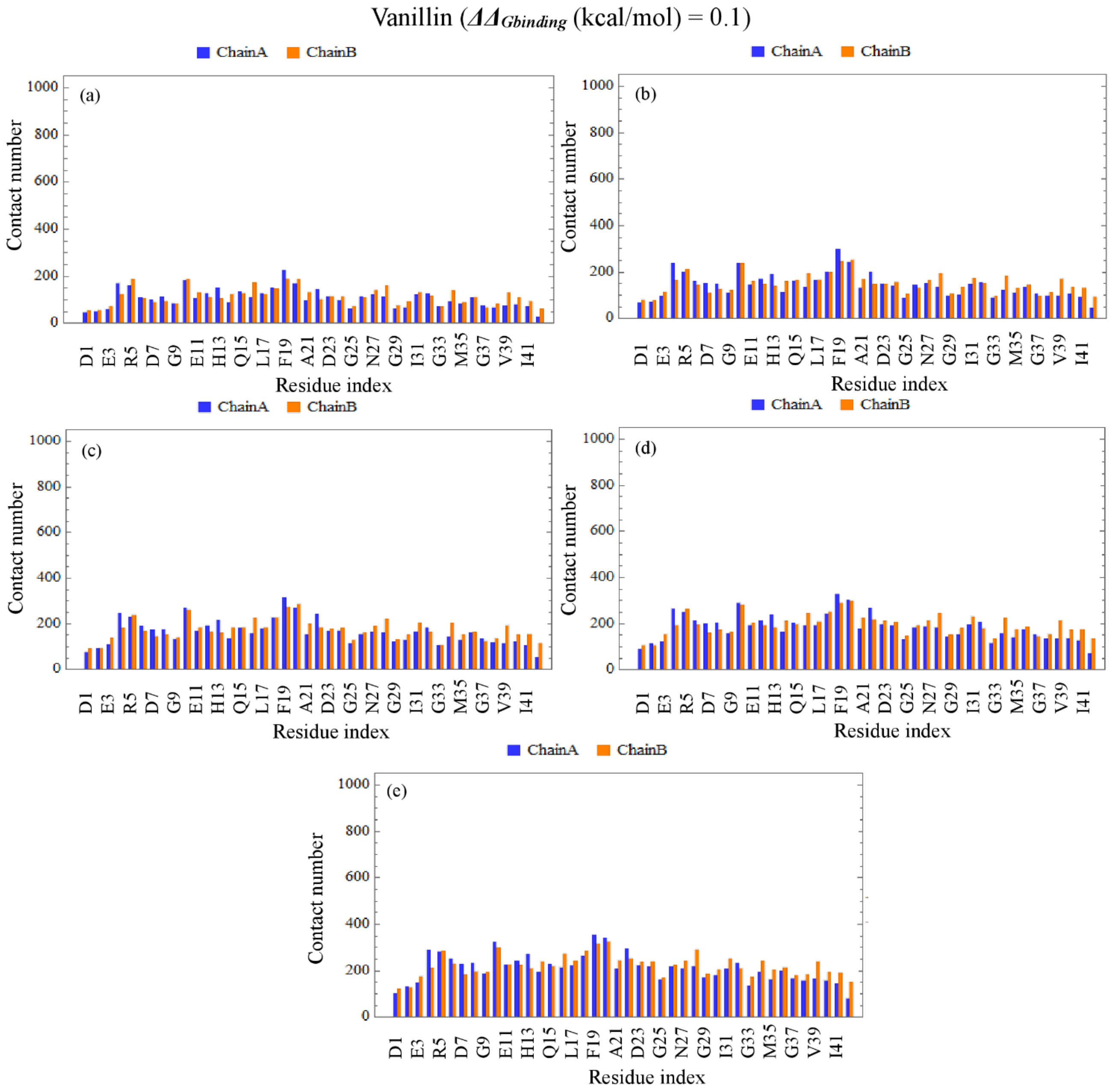


**Figure S10.** The number of contacts between individual residues and vanillin for $\Delta\Delta G_{binding}$ (kcal/mol) = 0.1 with five different distance cutoffs; (a) 0.3 nm, (b) 0.35 nm, (c) 0.4 nm, (d) 0.45 nm, and (e) 0.5 nm.

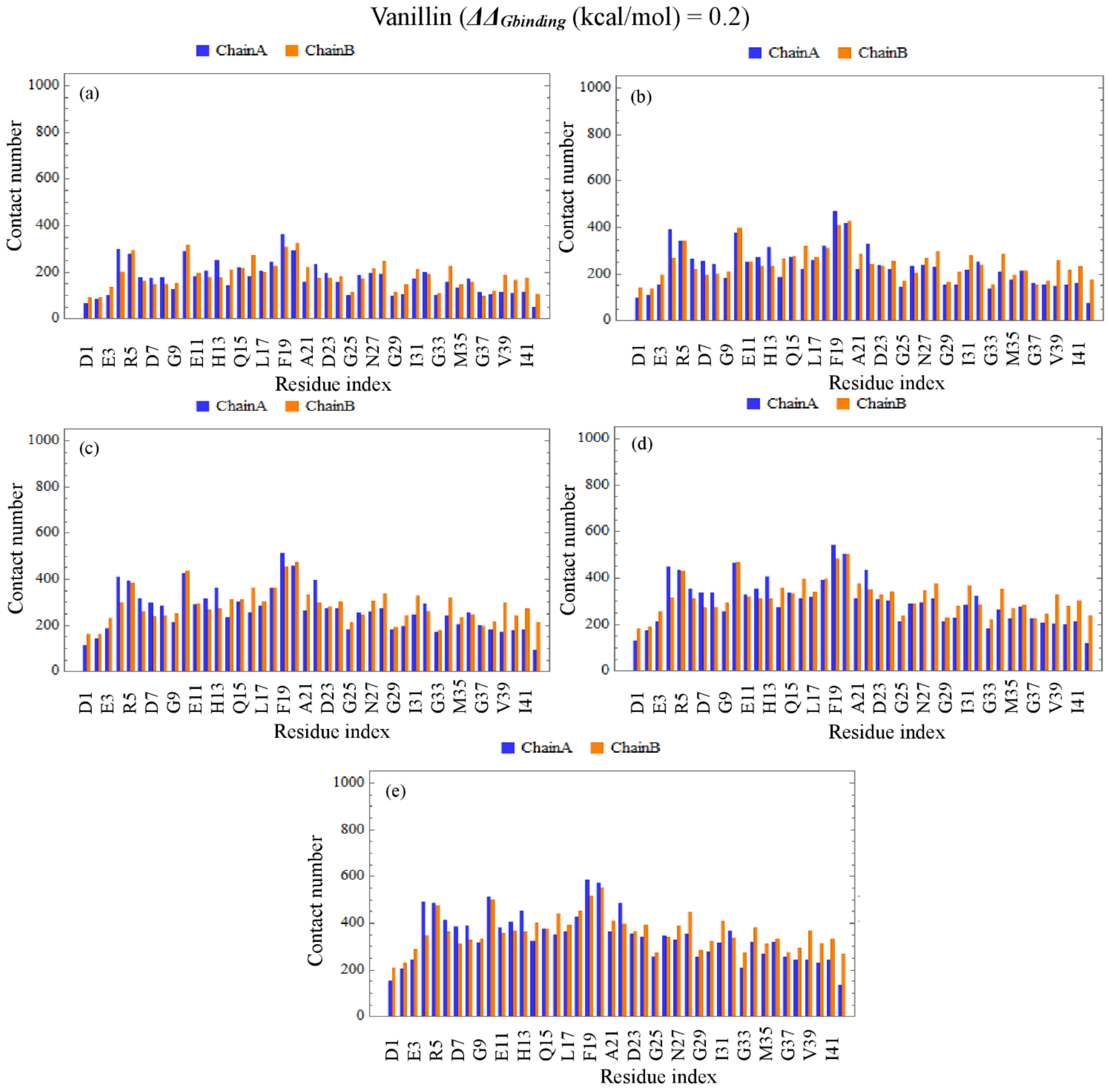


**Figure S11.** The number of contacts between individual residues and vanillin for $\Delta\Delta G_{binding}$ (kcal/mol) = 0.2 with five different distance cutoffs; (a) 0.3 nm, (b) 0.35 nm, (c) 0.4 nm, (d) 0.45 nm, and (e) 0.5 nm.

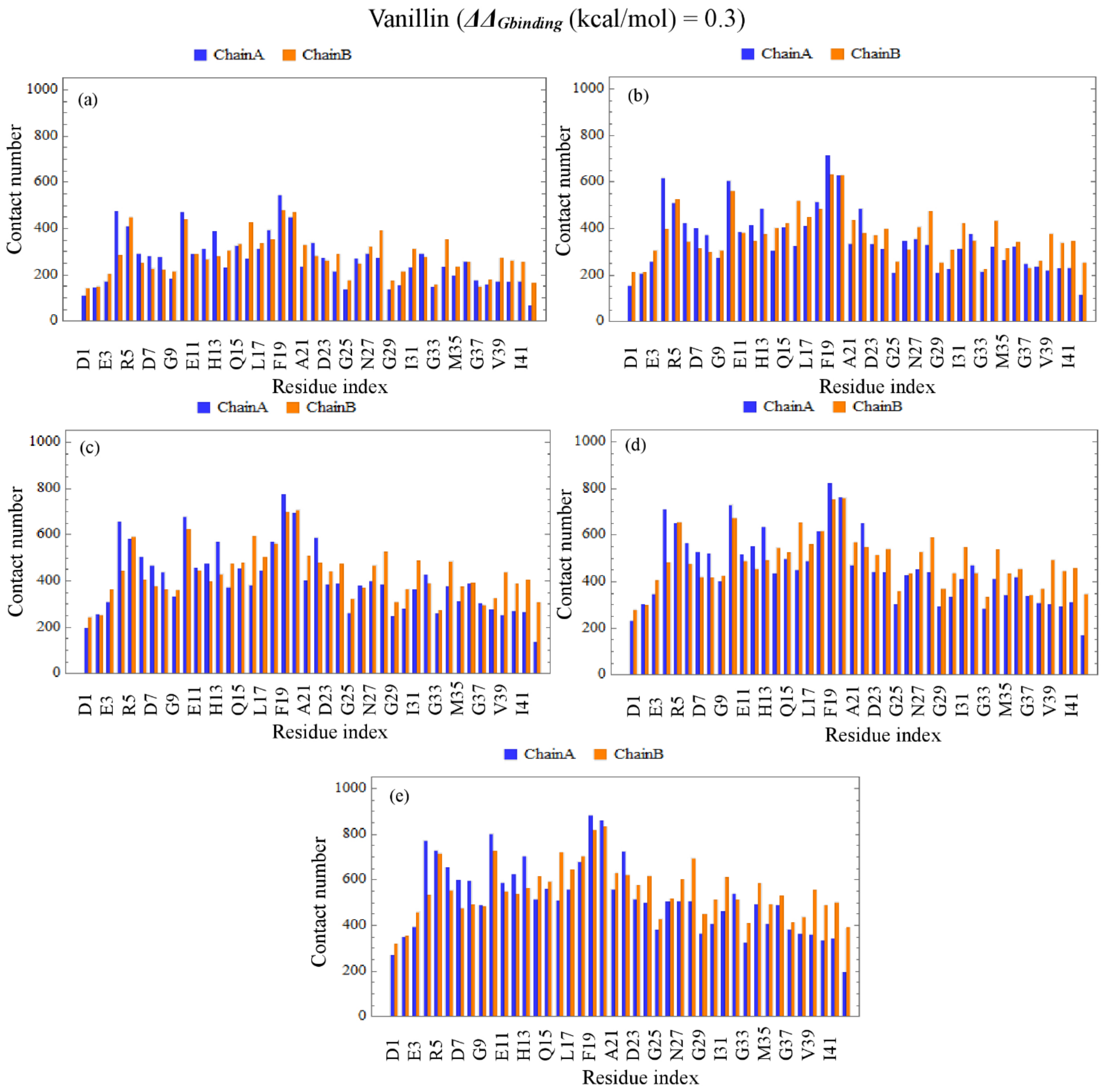


**Figure S12.** The number of contacts between individual residues and vanillin for $\Delta\Delta G_{binding}$ (kcal/mol) = 0.3 with five different distance cutoffs; (a) 0.3 nm, (b) 0.35 nm, (c) 0.4 nm, (d) 0.45 nm, and (e) 0.5 nm.
